# Population level rhythms in human skin: implications for circadian medicine

**DOI:** 10.1101/301820

**Authors:** Gang Wu, Marc D. Ruben, Robert E. Schmidt, Lauren J. Francey, David F. Smith, Ron C. Anafi, Jacob J. Hughey, Ryan Tasseff, Joseph D. Sherrill, John E. Oblong, Kevin J. Mills, John B. Hogenesch

## Abstract

Skin is the largest organ in the body and serves important barrier, regulatory, and sensory functions. Like other tissues, skin is subject to temporal fluctuations in physiological responses under both homeostatic and stressed states. To gain insight into these fluctuations, we investigated the role of the circadian clock in the transcriptional regulation of epidermis using a hybrid experimental design, where a limited set of human subjects (n=20) were sampled throughout the 24 h cycle and a larger population (n=219) were sampled once. By looking at pairwise correlations of core clock genes in 298 skin samples, we found a robust circadian oscillator in skin at the population level. Encouraged by this, we used CYCLOPS to reconstruct the temporal order of all samples and identified hundreds of rhythmically-expressed genes at the population level in human skin. We compared these results with published time-series skin data from mouse and show strong concordance in circadian phase across species for both transcripts and pathways. Further, like blood, skin is readily accessible and a potential source of biomarkers. Using ZeitZeiger, we identified a biomarker set for human skin that is capable of reporting circadian phase to within 3 h from a single sample. In summary, we show rhythms in human skin that persist at the population scale and a path to develop robust single-sample circadian biomarkers.

**One Sentence Summary:** Human epidermis shows strong circadian rhythms at the population scale and provides a better source for developing robust, single-sample circadian phase biomarkers than human blood.

## Introduction

Skin is the largest organ in the body, functioning in biosynthesis of vitamins and protection from the environment *(1, 2).* Daily variation in stressors (e.g., temperature, humidity, ultraviolet light) are met with corresponding changes in skin physiology. For example, blood flow, barrier recovery rate, temperature, cell proliferation, DNA repair, and transepidermal water loss all undergo diurnal variation *(3–6).* Importantly, core clock genes and hundreds of clock output genes show rhythmic or time-dependent expression in mouse and human skin *(7–9).* These rhythms are significant for skin health, e.g., to protect and repair daily UV radiation damage. In addition, disrupted circadian rhythms are associated with skin diseases, like psoriasis and skin cancer *(10, 11).* Yet our knowledge of circadian biology and the impact of photoaging on these processes in human skin is limited.

In addition to better understanding the circadian system in human skin, it also represents a potentially important source of circadian biomarkers. For circadian biology to influence healthcare, e.g. when to take a drug *(12)* or have a procedure *(13)*, a practical measure of circadian phase is necessary. The current ‘gold standard’ tool for assessing human circadian phase is dim light melatonin onset (DLMO). This assay requires a subject to sit in a dim room for repeated saliva sample collection; a difficult practice to standardize and perform at scale. Instead, most circadian biomarker research has focused on human blood transcriptomics with limited clinical success *(14, 15).* However, the transcriptional clock in individual human blood cells has both a lower amplitude and signal to noise ratio than the clock in individual fibroblasts *(16).* Moreover, the composition of blood varies greatly over time and across subjects due to internal (endocrine, body temperature, individual variation) and external factors (diet, infection). Together, these issues raise the question, is blood is the best source material for circadian biomarkers?

Here, we explore the potential for an alternative tissue, skin, to serve as a robust marker of human circadian phase. We describe a hybrid experimental design, where a limited set of subjects (n=20) were sampled throughout the 24 h cycle and a larger population (n=219) were sampled once. Using a recently described algorithm, cyclic ordering by periodic structure (CYCLOPS) *(17)*, we reconstructed the circadian transcriptome from human skin, finding hundreds of clock-regulated genes whose rhythms persist at the population level. We compared these results with time series data collected from mouse skin and show strong concordance in circadian phase across species at the transcript and pathway level. Finally, we applied a second algorithm, ZeitZeiger, to these reconstructed time course data. We report a biomarker set for human skin that is capable of reporting circadian phase to within 3 h from a single sample. In summary, we present a population scale survey of circadian rhythms in human skin and point to skin as a source of robust single-sample circadian biomarkers.

## Results

### Clock regulated genes identified from time series analysis of human skin

To explore the potential of human skin as a source of circadian phase biomarkers, we performed a time-series analysis of human skin every 6 h accrossa 24 h sampling period from 19 individuals (one subject with 3 samples was excluded). Each subject had a 2 mm full-thickness punch biopsy collected from their volar forearms every 6 h starting at 12 PM for a total of 4 biopsies. We used MetaCycle’s meta3d function to analyze these data. We identified 233 genes (*P* < 0.1; fig. S1A) that varied with a rhythmic pattern over a 24 h time period. These genes show a bimodal distribution, with peak phases clustered at 8-9 AM and 8-9 PM (fig. S1B). Phase set enrichment analysis (PSEA) *(18)* of these genes identifies L1 Cell Adhesion Molecule (L1CAM) interactions, signaling by NGF, immune system, metabolism of amino acids and derivatives, and metabolism of lipids and lipoproteins (fig. S1C) as pathways significantly enriched for cycling.

We reasoned that genes whose circadian pattern showed evolutionary conservation between mice and humans would represent a robust source for circadian biomarkers. If it’s conserved across 100 Mya of evolution, it should be conserved across different populations of humans. To identify these genes using a common statistical framework, we applied MetaCycle’s meta2d function to a recently published time-series dataset of mouse telogen (containing small and resting hair follicles) and anagen skin (containing large and growing hair follicles) *(7).* Our re-analysis found 1,280 and 294 circadian genes in telogen and anagen skin, respectively (*P* < 0.05). We noticed a pronounced bimodal phase distribution from mouse telogen, similar to human, and as previously noted from analysis of many other mouse tissues *(19)* (fig. S1D). Expression patterns for seven genes (*ARNTL*, *NPAS2*, *NR1D2*, *PER3*, *HLF*, *PER2* and *FKBP5*) were conserved across all three datasets (Fig. 1 and fig. S1E). We compared the expression profiles of each of these 7 genes across all human subjects. Although there were clear inter-individual differences (e.g., subjects 116 and 119, which show delayed melatonin and cortisol phase. fig. S2), the patterns of expression for these 7 genes generally followed the same phase relationship (Fig. 1). These genes were also robustly rhythmic in mouse telogen and anagen, albeit, with a nocturnal pattern of expression (Fig. 1). As previously noted, the phase of clock gene expression reflects locomotor activity, i.e. *ARNTL* peaks at 8 to 9 PM in humans and ZT23 in mouse, predicting the sleep phase in both species. The phase of clock gene expression matches their established phase relationships: *ARNTL* precedes *NR1D2*, which precedes *PER2 (20).*

**Fig. 1.**
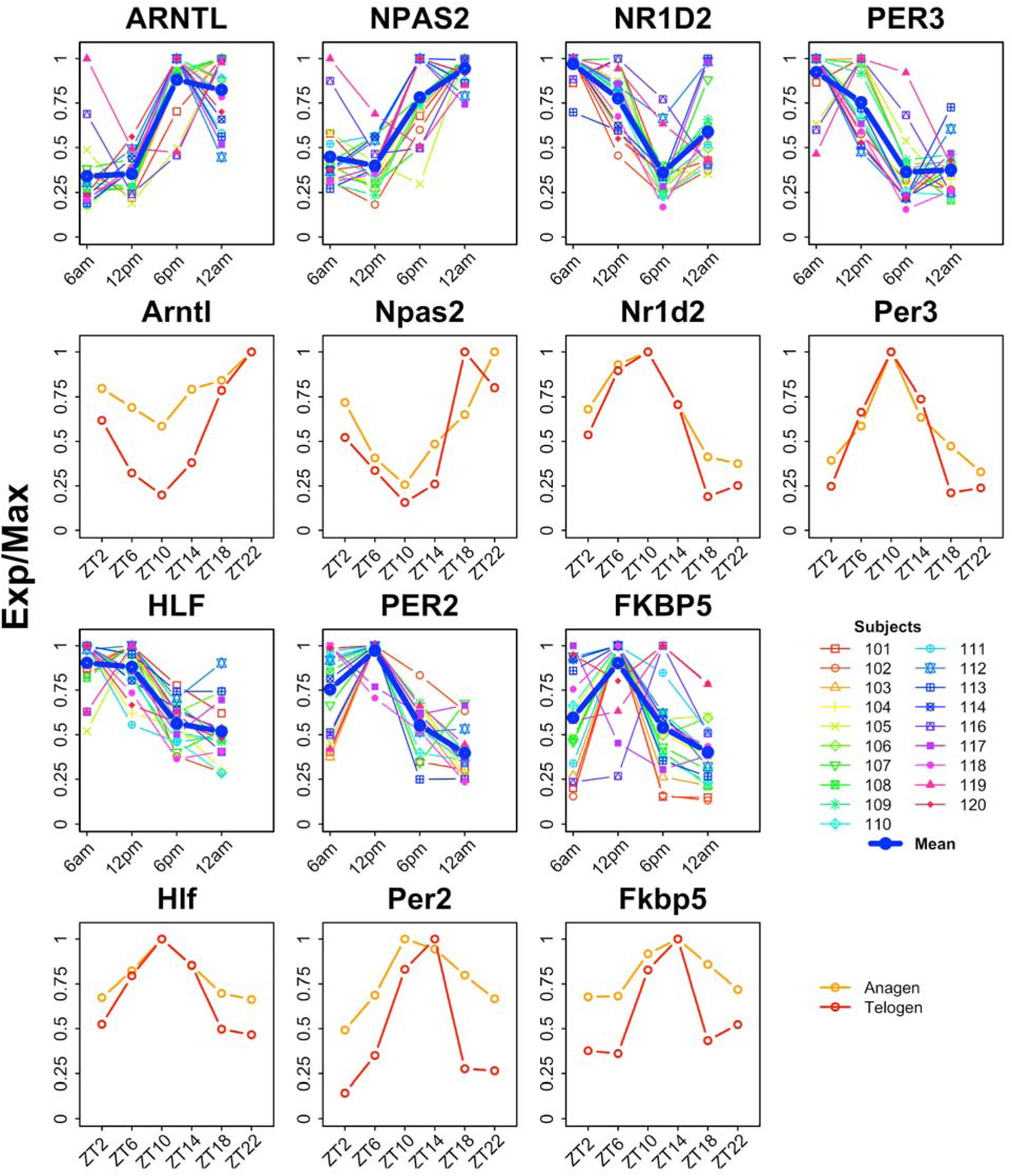
Individual variation of 7 conserved circadian genes. Skin from volar forearms of human subjects (n = 19) was sampled every 6 h for 1 circadian cycle starting at 12 PM. Expression profiles from all 19 human subjects along with the mean (blue) are shown (rows 1 and 3). Expression profiles for mouse anagen (orange) and telogen (red) skin samples are shown (rows 2 and 4). Mouse data are from *(7)* (table S1). Exp/Max indicates the expression value at each time point was normalized by the maximum expression across multiple time points in different human subjects or in mouse anagen or telogen tissue.

### Reproducing human skin phase without wall time

For skin to be a good source for determining human circadian phase, clock gene phase relationships should be conserved at a population scale. In other words, the circadian variation in clock gene expression must exceed inter-individual variation. We analyzed a gene expression dataset from human skin from 19 subjects sampled around a 24 h clock, and 219 subjects each sampled once irrespective of wall time (details in table S1). This hybrid design (fig. S3) captures advantages of both longitudinal and population based studies, while mitigating disadvantages. Using a set of core clock and strong output genes *(21)*, we used Spearman’s rho to evaluate the correlation of each gene against all others *(22)* (Fig. 2A). If the clock is intact, *ARNTL* should correlate positively with its partner transcriptional activator *NPAS2* and negatively with its target *NR1D1*, which represses *ARNTL.* As expected, we saw strong positive correlations among retinoid-related orphan receptor (ROR) targets *ARNTL*, *NPAS2*, *CLOCK* and strong negative correlations between these genes and their E-box targets *DBP*, *NR1D1*, *PER3* (Fig. 2, A and B). To reconstruct the daily rhythm of genome-wide gene expression in human skin, we applied CYCLOPS (see Materials and Methods). First, we evaluated the accuracy of the reconstructed sample order by comparing clock gene phase order between humans and mice (Fig. 2C). By both visual inspection and statistical analysis (Fisher’s circular correlation = 0.715), the reconstruction of the human circadian cycle (Fig. 2C, outer circle) matched the mouse (Fig. 2C, inner circle). In addition, we compared CYCLOPS-predicted sample phase for each of the 19 subjects for which sampling time was available. The strong linear relationship between the mean CYCLOPS-predicted phase and sampling times suggests accurate sample ordering (Fisher’s circular correlation = 0.953; Fig. 2D).

**Fig. 2.**
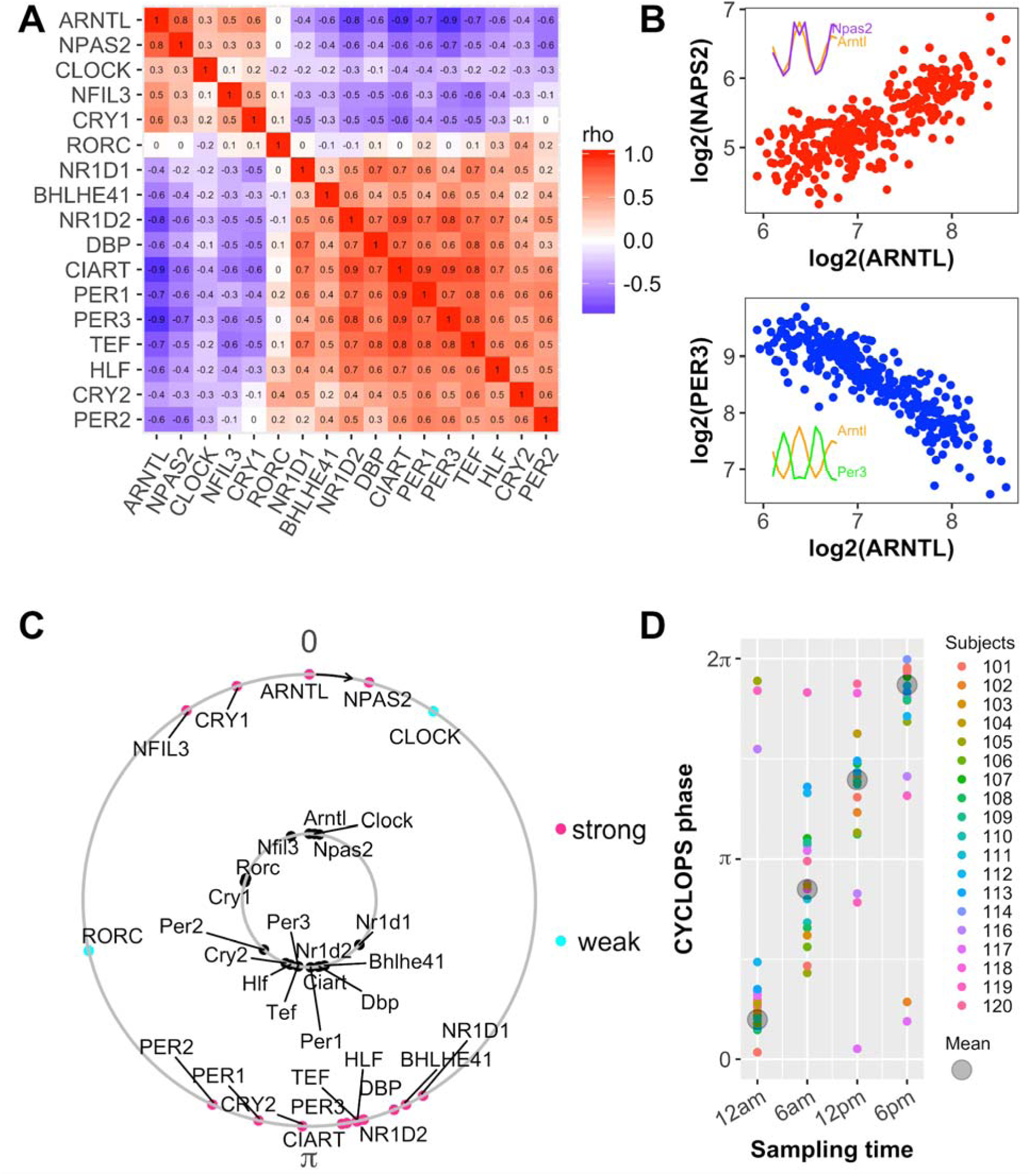
Evaluation of circadian function in human skin. (A) A heat map of Spearman’s rho for core clock and output genes from ordered data (n = 20, table S1) and unordered data (n = 219, table S1) shows conserved correlation structure of clock genes. (B) Examples of correlated clock gene expression (*ARNTL* and *NPAS2*, red) and anti-correlated clock gene expression (*ARNTL* and *PER3*, blue). Each point represents one human skin sample. Expression profiles of corresponding mouse clock genes from telogen (orange, purple and green are shown for *Arntl*, *Npas2* and *Per3*, respectively). (C) CYCLOPS was used to order human data from all 239 subjects. In an intact clock, ROR phased genes (e.g. *ARNTL*, *NPAS2*, *CLOCK*) always peak prior to E-box phased genes (e.g. *NR1D1*, *DBP*, *PER1*). Conserved phase relationships are shown for clock-regulated genes (internal circle, mouse; external circle, human). For human genes, pink shows strong cycling (FDR < 0.15, rAMP > 0.1, rsq > 0.1), while cyan shows weaker cycling. The phase of *Arntl* (mouse) or *ARNTL* (human) is set as 0 to facilitate comparison. (D) CYCLOPS accurately recalls circadian phase from 19 subjects, 4 timepoints/subject. Different colors indicate different subjects, and the circular average phases for all samples is shown in grey.

### Human transcriptional rhythms are evolutionarily conserved

We expanded our analysis of rhythms in human skin to the broader transcriptome from 298 samples collected from 238 donors. Using the CYCLOPS modified cosinor regression algorithm, we selected 135 circadian genes (false discovery rate (FDR) < 0.15, relative amplitude (rAmp) > 0.1, goodness-of-fit (rsq) > 0.1, and fitmean > 16) (Fig. 3A, fig. S4 and Data file S1). While ROR/REV-ERB-response element (RORE) and E-box phases had dominant signatures, other phases were also represented (Fig. 3A). Our central hypothesis is that high amplitude oscillatory genes with conserved phase relationships across species will be the most robust biomarkers. Comparing periodic genes from mouse telogen, we defined five categories of genes (Fig. 3B), those with i) no mouse homologs, ii) low amplitude and no significant cycling in mouse, iii) high amplitude and no significant cycling, iv) low amplitude and cycling, and v) high amplitude and cycling. More than half of the rhythmic genes in humans (n = 96) were not significant in mouse telogen (Fig. 3B). For those that were robustly cycling in mouse telogen, we compared their phases and found a strong linear correlation for most genes (Fig. 3C). We investigated the underlying biology using PSEA and identified temporal regulation of nine significantly enriched pathways at the population level (fig. S5A). Five of these pathways are also enriched in mouse telogen skin (fig. S5, B and C), including pathways related to cell cycle, adaptive immune system, immune system, matrisome, and circadian clock. Interestingly, in addition to the pathways themselves, their phase order is conserved between mice and humans (fig. S5C), demonstrating the conserved orderly progression of skins gene expression program.

**Fig. 3.**
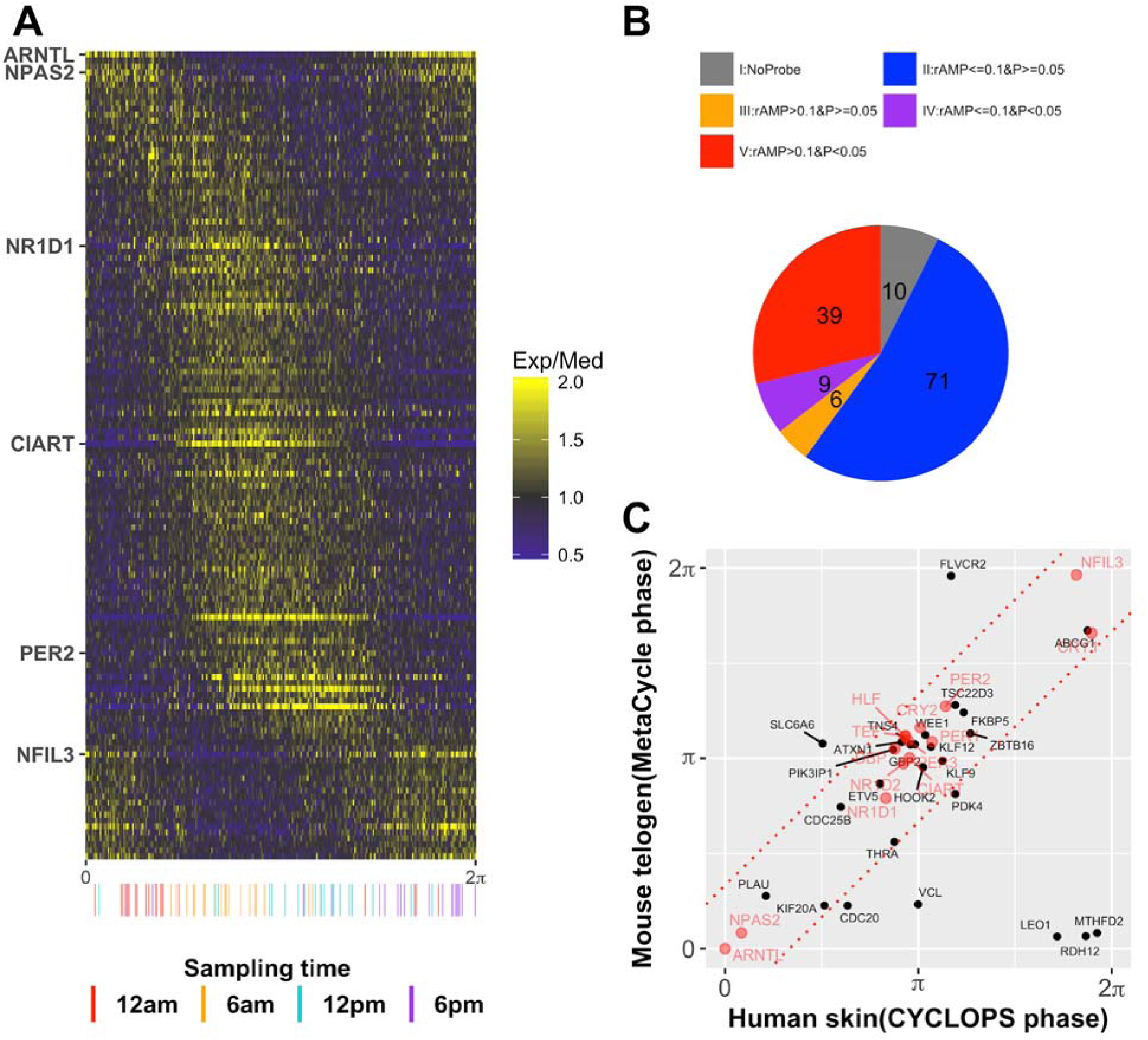
Population analysis of human skin data reveals conserved circadian transcriptional output. (A) CYCLOPS was used to phase-order each sample from 239 subjects, followed by modified cosinor regression to identify clock regulated genes. A heatmap of median centered gene expression values for all human samples ordered by phase and all significant clock regulated genes (n = 135) is shown. Clock genes that best depict circadian phases are labeled on the Y axis (e.g. *ARNTL*, *NR1D1*). Samples with known sample acquisition times (n = 79) are on the X axis. (B) A pie chart of all human circadian genes and expression of their homologs in mouse telogen. Groups are: no mouse homolog (grey), low amplitude (rAMP ≤ 0.1) and not cycling (*P* value ≥ 0.05) (blue), high amplitude (rAMP > 0.1) and not cycling (orange), low amplitude and cycling (*P* value < 0.05) (purple), and high amplitude and cycling (red). (C) A comparison of high amplitude cycling genes (Group V, n = 39 genes) reveals strong phase conservation between mice and humans. Core clock genes are shown in red. The predicted gene phases from CYCLOPS and MetaCycle were adjusted to *ARNTL*/*Arntl* phase (set to 0).

### Population-level biomarkers of circadian phase in human skin

In addition to analyzing clock-regulated biology, these datasets present an opportunity to discover robust biomarkers of circadian phase that persist at the population level. To date, studies performed on human blood suggest high variability and low amplitude rhythms from mixed cell types *(23).* Furthermore, when Brown and colleagues analyzed single cell type from blood, they found the cell autonomous oscillator in blood monocytes was significantly weaker than in fibroblasts *(16).* To clarify which human tissue is a better source of circadian biomarkers, we compared the expression correlation matrices of 10 core clock genes *(24)* based on time-series samples from human blood (n = 201) *(25)*, skin (n = 79) and multiple mouse tissues (n = 144) *(19).* The co-expression pattern of these 10 core clock genes across 12 mouse tissues is quite similar (Mantel test, *P* = 3e-06) with human skin, but is not similar with human blood (Mantel test, *P* = 0.897; Fig. 4A). At population level, the co-expression pattern of clock genes in skin (Fig. 2A) is also stronger than in isolated monocytes or T cells collected from hundreds of individuals (fig. S6). This is in agreement with prior studies and suggests that skin is a better tissue for identifying robust circadian biomarkers. We used a recently published method, ZeitZeiger *(26)*, to identify biomarkers that could accurately report circadian phase. ZeitZeiger was trained with CYCLOPS assigned phase from all 219 samples without sampling time and 43 samples with sampling time. This resulted in a set of 29 candidate biomarkers (Fig. 4B), approximately half expressed in two different phases, represented by two sparse principal components (Fig. 4B, SPC1 and SPC2). To validate these biomarkers, we analyzed a set of 9 human subjects each measured at 4 separate times of day. The average predicted phase was within 3 h at all four points comparing with the anticipated time (Fig. 4C). For 6 of the subjects, it was difficult to distinguish between 6 PM and 12 AM samples. When we evaluated the sample relationships for different times of day, the median absolute error was 2.95 h for the 9 subjects in this study (fig. S7). Nevertheless, we were able to accurately assign 30/36 of the samples to the correct circadian phase.

**Fig. 4.**
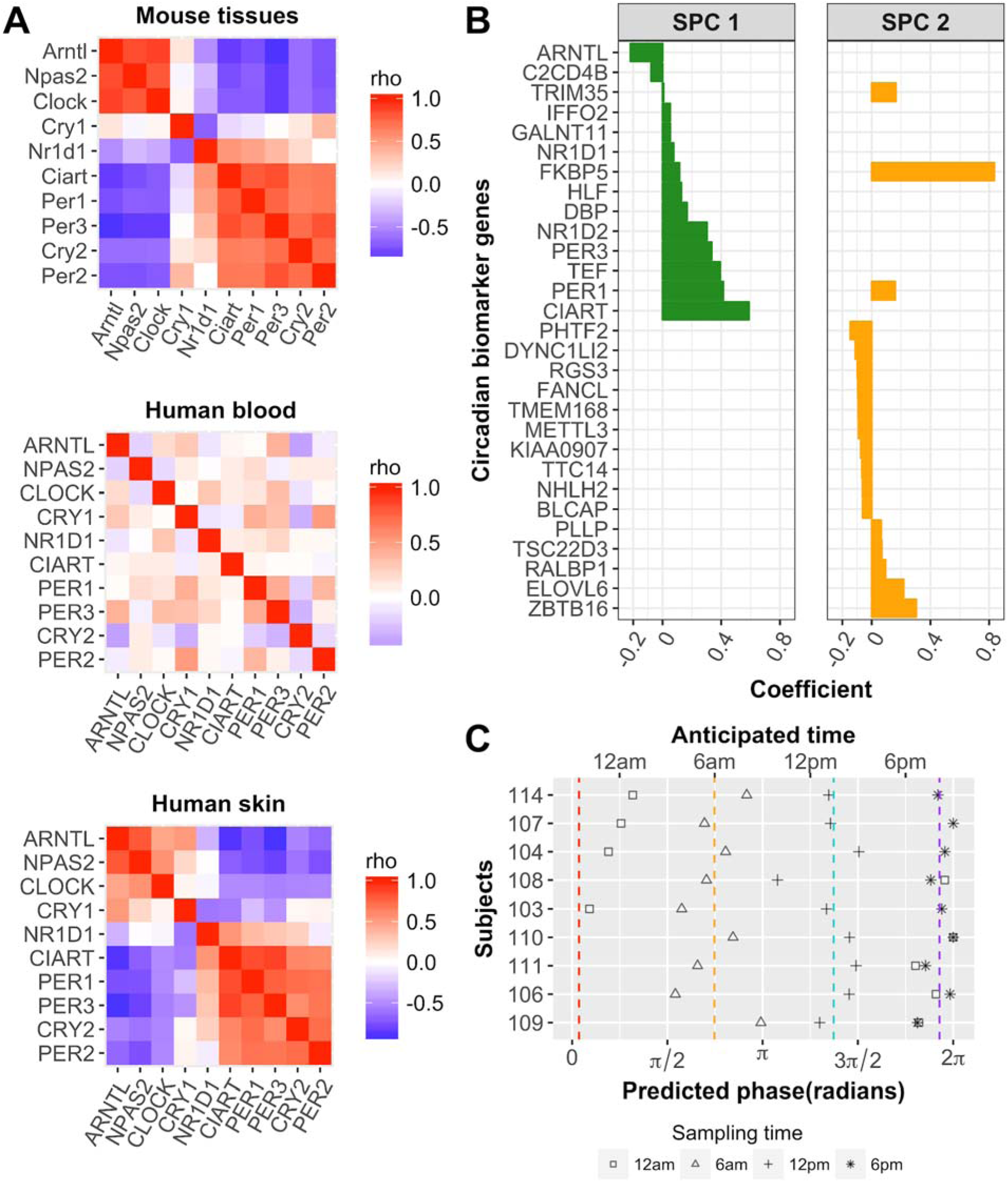
Population level biomarkers of circadian phase from a single human skin sample. (A) Evaluation of circadian clock function from 12 mouse tissues (top panel), human blood (middle), and human skin (bottom). The mouse data are from Zhang and Lahens *(19)* and are included as a benchmark. The clock in human blood is much weaker than in human skin. Red and blue indicate positive and negative Spearman’s rho values, respectively. (B) ZeitZeiger *(26)* was used to select a set of 29 circadian marker gene candidates with absolute coefficient values greater than 0.05, as described in Supplementary Materials and Methods. (C) Validation of predicted markers. Using the circadian marker set and ZeitZeiger, we analyzed samples collected every 6 h over a circadian day for 9 subjects that were excluded from the training set. Average predicted phases of samples collected at 12 AM, 6 AM, 12 PM and 6 PM are indicated with red, orange, cyan and purple dashed line, respectively. Sample phase prediction around the E-box phase (orange, cyan, and purple) was generally accurate, while sample phase prediction between ROR and E-box phase (red) did not perform as well.

## Discussion

Here we report a population-based analysis of circadian rhythms in human skin. We used a hybrid experimental design (fig. S3) to examine circadian gene expression, where 20 subjects were sampled longitudinally and 219 were sampled once without regard to circadian time. We show that clock gene correlations are strong and evident in unordered data from human skin, unlike human blood. We used our recently published algorithm, CYCLOPS *(17)*, to order these samples and explore population level transcriptional rhythms. As predicted, the phases of core clock machinery and most clock outputs are shared between mice and humans. For pathways that are clock regulated in both species, the phase order is preserved with the cell cycle peaking early in the inactive phase, followed by the immune system, matrisome, and the E-box regulated genes of the circadian clock. Finally, prompted by the observation that human skin appears to be highly rhythmic, we used a recently published algorithm, ZeitZeiger, to look for a gene set that accurately reports circadian phase from a single sample. These results have broad implications for the core clock, skin physiology, and circadian medicine.

Core clock genes tend to oscillate in most, if not all, organs. We found that *FKBP5*, like core clock genes, oscillates in both mouse skin and also at the population level in humans (Fig. 1 and Fig. 3C). *FKBP5* is an immunophilin protein family member involved in protein folding and trafficking that is thought to mediate calcineurin inhibition *(27)*; it cycles in seven additional mouse tissues (liver, kidney, aorta, heart, distal colon, lung and pituitary; JTK q-value < 0.05) *(28).* Taken together, these data suggest that *FKBP5* may represent a previously unrecognized component of the clock. Notably, the calcineurin inhibitor tacrolimus is a topical anti-inflammatory therapeutic for the treatment of atopic dermatitis *(29).* Given our findings, it stands to reason that time of day could be leveraged to optimize drug efficacy for this and other skin conditions.

We also found that several components of the cell cycle machinery robustly cycle in human skin. These include *CDC20*, *CDC25B*, *KIF20A*, and *WEE1.* Their encoding proteins are all associated with mitosis (Fig. 3C and fig. S5C) and may contribute to circadian phase dependent proliferation of epidermal cells *(7, 30).* Interestingly, mouse liver cells regenerate in a *Wee1*-dependent fashion at a single phase in the circadian cycle *(31).* The cellular clock in human skin may also use *WEE1* to gate epidermal stem cell division. This process is key to epidermal homeostasis, as new cells replace those lost during turnover or injury *(32).*

Biomarkers capable of reporting circadian phase are essential for circadian medicine. The current standard in the field, DLMO, is impractical and burdensome. Many groups have sought circadian biomarkers from blood, including analysis of metabolites and gene expression *(14, 15, 33)*. However, blood is a heterogeneous mixture that changes composition in response to diet, infection, exercise, and many other factors, likely explaining the weak rhythmic expression of clock genes in blood monocytes *(16)* and time-series blood samples *(23).* Relatively little attention has been paid to other accessible human sources, such as skin.

Hypothesizing that phase relationships of clock genes will be preserved in unordered data, we used an algorithm *(22)* to directly compare human skin and blood. Strikingly, we found the oscillator in unordered data from human skin (Fig. 2A) is far stronger than in isolated monocytes or T cells collected from hundreds of individuals (fig. S6). Unlike blood, clock genes are robustly cycling in mammalian skin *(7, 8, 34)* and the expression relationship of clock genes is conserved across species (Fig. 4A). Further, the clock in skin is controlled by the SCN *(34)* and can be shifted by time-restricted feeding *(35)*, which suggests circadian phase of internal organs (e.g. gut, liver) may be reflected in the skin transcriptome. Taken together, these findings suggest human skin has a stronger oscillator that is clearly evident at the population-scale in humans. We suggest that circadian biomarker discovery should include a focus on human skin.

Our study has limitations. For example, the relatively few samples (~50) collected during the night phase limit statistical power. In addition, the majority of samples were taken from healthy females, perhaps limiting applicability to broader patient populations. To overcome these limitations, future experiments should include targeted sampling of the inactive phase and control for other demographics. Clinical use of circadian biomarkers will require fast, cost-effective, and non-invasive sampling techniques (e.g. tape strip based, hair follicle or oral mucosa based) *(36)* and standardization across platforms (e.g. qPCR or array). Larger studies would also benefit from a comparison to other standardized techniques (e.g. DLMO, temperature, and actigraphy). Finally, consistency of results should be confirmed across a range of disease states and pathologies.

Our data showing strong, population level rhythms in human skin, however, suggests that development of single-sample circadian biomarkers is possible. Recent research shows that the timing of medical intervention can impact patient response *(13)* and reviewed in *(12).* The identification of a biomarker set that accurately predicts systemic circadian phase to within 3 hours from a single sample is a major step towards translating circadian biology to medicine.

## Materials and Methods

### Human clinical experimental design and sample collection

Both clinical studies adhered to the International Council on Harmonization of Good Clinical Practices, and the principles expressed in the Declaration of Helsinki. Associated protocols were approved by Institutional Review Boards, and each subject provided informed consent. For collecting the ordered samples, 20 healthy Caucasian male subjects, aged 21-49; median age 24.5, free of skin disease and with a normal sleeping pattern (6-8 hr/night) were recruited and housed in a facility that specializes in sleep studies (Community Research, Cincinnati, OH). Saliva samples were collected every three hours over the same 24 hour period to confirm normal cycling of cortisol and melatonin (fig. S8 and S9). Full thickness (2 mm) punch biopsies were collected from sites on the volar forearms of each subject at six hour intervals over a 24 hour period (6:00 A.M., 12:00 P.M., 6:00 P.M., 12:00 A.M.). Biopsies were processed for separation into epidermal and dermal compartments by Laser Capture Microdissection (LCM). Detailed protocol of LCM is included in Supplementary Materials and Methods. The collection of the unordered 152 Caucasian and 67 African-American female samples has been previously described *(37).*

### mRNA Target Labeling and Processing

Total RNA from LCM samples was isolated utilizing the Pico Pure RNA Isolation Kit (Life Technologies, Grand Island, NY) according to manufacturer’s recommendations. Quality and concentration of the isolated RNA was determined utilizing an Agilent 2100 Bioanalyzer and the RNA 6000 Pico Kit (Agilent Technologies, Santa Clara, CA) according to manufacturer’s recommendations. In brief, 25 ng of total RNA was reverse-transcribed into cDNA copies using oligo-dT primers and reverse transcriptase followed by second strand synthesis using DNA polymerase I. In vitro transcription synthesizes cRNA that is biotin incorporated and purified using the Affymetrix HT 3’ IVT Express kit (Cat. #901253), as executed on a Beckman Biomek FXp Laboratory Automation Workstation (Beckman Cat. #A31842). Biotinylated cRNA was fragmented by limited alkaline hydrolysis and then hybridized overnight to Affymetrix GeneTitan U219 array plates using the Affymetrix GeneTitan instrument.

### Time-series analysis with MetaCycle

The RMA algorithm *(38)* was used to extract expression profiles from the raw CEL files. MetaCycle::meta3d *(39)* and MetaCycle::meta2d was used to detect circadian transcripts from time-series expression profiles of human skin samples, and mouse anagen and telogen samples (4h-interval covering two days) *(7)*, respectively. A representative transcript with the smallest *P* value or the largest rAMP if multiple transcripts have equal *P* value was selected for each gene. Those genes meeting with given significance cut-off (*P* < 0.1 for human skin and *P* < 0.05 for mouse skin) and amplitude cut-off (rAMP > 0.1) were selected as circadian genes. Detailed parameter settings for running MetaCycle are included in the Supplementary Materials and Methods. Those circadian genes that overlapped in at least two of three datasets were used as seed gene list for CYCLOPS ordering.

### Ordering human skin samples with CYCLOPS

All skin samples (298 in total) were ordered with CYCLOPS (Julia version 0.3.12) *(17).* The fidelity of clock output was evaluated for 17 clock-controlled genes (CCGs), using a similar method to Shilts et al. *(22).* While doing this research, eigengene selection became limiting. To overcome this hurdle, we incorporated features of a second algorithm Oscope *(40)* to filter periodic out-of-phase eigengenes. We used seed genes selected from time series data to generate 10 eigengene clusters, which were evaluated by Oscope. CYCLOPS produced sample ordering based on relationship between multiple eigengenes. Orderings were evaluated by comparing phases of gene expression of 17 CCGs with their mouse homologs, and the best was selected by visual inspection and Fisher’s circular correlation coefficient. Using this ordering, modified cosinor regression was applied to each expressed gene. Genes with FDR < 0.15, fitmean > 16, rAMP > 0.1, rsq > 0.1 were designated as circadian. Additional details on the implementation of CYCLOPS and Oscope are described in the Supplementary Materials and Methods.

### Explore skin biomarkers with ZeitZeiger

ZeitZeiger *(26)* was used to explore the circadian marker genes with CYCLOPS ordered human skin samples. All skin samples were divided into two data sets — training and testing. The training set includes all 219 samples without sampling time and 43 samples with sampling time information collected from 11 subjects (each subject has four samples except subject 115). The testing set includes 9 subjects, selected based on expression profiles of circadian genes (Fig. 1). Circadian biomarkers were selected from the training dataset and used for predicting phase of the test set, as detailed in the Supplementary Materials and Methods.

### Statistical analysis

The following statistical methods were used in this study: Fisher’s method in MetaCycle, Kuiper’s test in PSEA, circular node autoencoders, modified cosinor regression, and the F test as implemented in CYCLOPS, K-medoids clustering in Oscope, maximum-likelihood estimation in ZeitZeiger, Spearman’s rank correlation, Mantel test, and permutation testing to compare the expression correlation matrix of core clock genes. Depending on the experimental design and statistical method, *P* value cutoffs of < 0.1 and < 0.05 were used for time-series analysis. *P* values from CYCLOPS were adjusted by the Benjamini-Hochberg procedure. Genes with FDR of < 0.15 were designated as rhythmically-expressed at human population level. Additional criteria are listed in the Supplementary Material and Methods.

## Supplementary Materials

Materials and Methods

Fig. S1. Comparison of clock-regulated genes identified from time-series analyses of human and mouse skin.

Fig. S2. Salivary melatonin and cortisol levels and clock gene expression profiles in subject 114, 116 and 119.

Fig. S3. Pros and Cons of experimental designs for human circadian biology.

Fig. S4. Number of circadian genes identified in human skin at a series of FDR cut-offs.

Fig. S5. PSEA analysis of circadian genes identified in human skin and mouse telogen.

Fig. S6. Weak circadian clock in human population-level blood samples.

Fig. S7. Validation of skin circadian markers.

Fig. S8. Daily salivary melatonin levels for 20 subjects.

Fig. S9. Daily salivary cortisol levels for 20 subjects.

Fig. S10. CYCLOPS recovers sample order from mouse anagen and telogen datasets.

Table S1. List of datasets used in this study.

Data file S1. List of genes cycling in human skin at population level.

## Acknowledgments

For the ordered data clinical study we thank Dr. Joseph Kaczvinsky and Kathleen Werchowski for clinical protocol development and Jim Li for statistical analysis. We also thank Drs. Bhavani Kasibhatla, Charles Bascom and Rosemarie Osborne for helpful discussions. We also thank Tiago de Andrade and Garrett Rogers for thoughtful discussions.

## Funding

Procter and Gamble paid for 100% of the costs of clinical work reported in this paper. This work is also supported by the National Institute of Neurological Disorders and Stroke (7R01NS054794-11 to JBH and Andrew Liu), the National Human Genome Research Institute (2R01HG005220-5 to Rafa Irizarry and JBH), and (1R21NS101983-01 to Tom Kilduff and JBH), the Defense Advanced Research Projects Agency (D17AP00003 to RCA) and the National Institute of General Medical Sciences (R35GM124685 to JJH).

## Author contributions

J.B.H., J.E.O., and K.J.M, designed research; G.W., M. D. R, R. E. S., R.T., J.D.S. and L. J. F. performed research; R. C. A., J. J. H. and J. B. H. contributed analytic tools; G.W., M. D. R. and R. E. S. analyzed data; J. B. H., G. W., M. D. R., L. J. F., D. F. S. wrote the paper.

## Competing interests

K.J.M., J.E.O., J.D.S. and R.T. are employees of The Procter and Gamble Company, which markets skin care products.

## Data and materials availability

All gene expression data have been deposited in GEO (GSE112660).

